# StretchfMRI: a new technique to quantify the contribution of the reticular formation to long-latency responses via fMRI

**DOI:** 10.1101/582692

**Authors:** Andrea Zonnino, Andria J. Farrens, David Ress, Fabrizio Sergi

## Abstract

Increased reticulospinal (RS) function has been observed to cause both positive and negative outcomes in the recovery of motor function after corticospinal lesions such as stroke. Current knowledge of RF function is limited by the lack of accurate, noninvasive methods for measuring RS function. Recent studies suggest that the RS tract may be involved in processing and generating Long Latency Responses (LLRs). LLRs, elicited by applying precisely controlled perturbations, can act as a reliable stimulus to measure LLR-related brainstem function using fMRI with high signal-to-noise ratio.

In this paper, we present StretchfMRI, a new technique that enables simultaneous recording of neural and muscular activity during motor responses conditioned by velocity-controlled robotic perturbations, which allows for direct investigation of the neural correlates of LLRs using fMRI.

Via preliminary validation experiments, we demonstrate that our technique can reliably elicit and identify LLRs in two wrist muscles–FCR and ECU. Moreover, via a single-subject pilot experiment, we show that the occurrence of an LLR in a flexor and extensor muscle modulates neural activity in distinct regions of the brainstem. The observed somatotopic organization is in agreement with the double reciprocal model of RS function observed in animal models, in which the right medullary and left pontine reticular formation are responsible for control of the motor activity in flexors and extensors, respectively.

## I. Introduction

Stroke is a leading cause of disability, affecting more than one new US citizen every minute, 50% of which are left with residual motor disability [1]. Motor disabilities are mainly a consequence of the reduced corticospinal drive that results in a loss of movement accuracy, in a decreased ability to independently control joint movement, and in muscles flaccidity. To this date, different rehabilitative interventions have been proposed to promote motor recovery based on either traditional or robotic [2] therapy. Both approaches aim to promote neuroplasticity with the ultimate goal to help the patients restore the natural corticospinal drive and re-use the same body segments in a way that resembles unimpaired conditions as closely as possible.

However, there are multiple other descending pathways that concur with the corticospinal tract in controlling generation of voluntary motor actions [3]. Within these secondary motor pathways, the reticulospinal tract (RST) is especially important as it has been shown to be involved in locomotion control [4], maintenance of posture [5], reaching [6] and grasping [7]. While previous research advanced that the RST could be a useful target for neuro-rehabilitative interventions that aimed to strengthen neural drive to skeletal muscles [8]–[10], recent research also shows that in humans RST function is associated with post-stroke impairment, including loss of independent joint control and hyperexcitability of stretch reflex [11], [12].

The RST consists of two different pathways that originate from the Reticular Formation (RF), a region in the brainstem composed of a constellation of multiple nuclei [13], [14], and connect to the motorneurons via segmental interneurons. Animal experiments advance a possible organization of motor function in the RF, where the different nuclei provide excitatory signals for ipsilateral flexors and contralateral extensors, while inhibitory signals are provided to contralateral flexors and ipsilateral extensors [10]. However, it is currently unknown if this model is accurate in humans, mostly because accurate imaging of RF function in humans is an open challenge.

Recent studies advanced the hypothesis that the RF may be involved in processing long-latency responses generated by feedback control of human movements [15]. Identified over 60 years ago [16], [17], the long-latency response (LLR) is a set of muscle activation bursts that, for the upper extremity, occur 50-100 ms after an undesired limb displacement stretches the spindles of a set of muscles. LLRs are a fundamental component of motor control, as they gracefully blend the fast reaction time afforded by reflexes with the flexible and skilled action of voluntary movement [18]. Thereby, investigating how the neural activity in the RF is modulated by the generation of a LLR would be a powerful tool to improve our understanding of the role of the RF in motor control. Moreover, as LLRs are “semi-reflexive” responses, confounds that usually affect voluntary motor tasks, such as individual subject skill and task-related performance, are expected to affect the neurophysiology of LLRs with smaller between-subject variability, opening the possibility to directly measure the activity in the RF using neuroimaging.

In this paper we present StretchfMRI, a novel technique that we have developed to study the neural substrates of LLR *in-vivo* in humans. StretchfMRI combines robotic perturbations with electromyography (EMG) with functional Magnetic Resonance Imaging (fMRI) allowing to obtain simultaneous recording of neural and muscular activity during the generation of LLRs. This paper focuses on our novel protocol detailing three main aspects: the design of a novel MR-compatible wrist robot, the perturbation protocol used during fMRI imaging, and the acquisition and processing of the EMG data. We also present a set of experiments conducted to validate the different components of our technique. Finally, the integrated StretchfMRI protocol was used to determine the neural substrates of LLR in a pilot, single-subject study.

## II. Material and Methods

Investigations of the neural substrates of LLR via fMRI requires simultaneous acquisition of muscular and neural activity during the amplication of velocity-controlled perturbations that elicit LLRs in a known set of upper extremity muscles. While previous studies primarily investigated the reflex activity of proximal muscles by applying perturbations to the shoulder and elbow joints [15], we focused our attention to forearm muscles during wrist perturbations. By targeting wrist movements, we expect to have reduced head and body motion, two factors that can degrade the quality of the fMRI images [19] and the EMG recordings.

### A. Simultaneous recording of fMRI and EMG data

Measuring EMG during fMRI protocols is technically challenging because of the large artifacts introduced in the EMG recordings by the coupling of the fMRI time- and spatially-varying electromagnetic fields–i.e. static magnetic field, gradient magnetic field, radio wave–and the undesired movement of the EMG electrodes. While the radio waves introduce noise at a frequency that is far from the spectrum expected for physiological muscle contractions, and so it a can be easily removed using standard low-pass filters, the same is not true for gradient and movement artifacts, whose spectrum overlaps with the one expected for muscle contractions.

Different algorithms have been proposed to compensate for the gradient artifacts in Electroencephalogram (EEG) and EMG protocols [20], [21], however, they are not capable of fully removing the artifacts when they couple with electrodes movement. Given our interest in only short periods of the EMG recordings–i.e. those where we expect to observe a LLR–we pursued a simplified approach to avoid gradient artifacts. We introduced in the MRI scanning protocol a 200 ms silent window after each acquisition volume where no fMRI excitation is generated. By synchronizing the application of the velocity controlled perturbations with the timing of the fMRI sequence it was then possible to obtain gradient artifact-free measurements of LLR (Fig. 1). Doing so, we leveraged the intrinsic delay of the hemodynamic signal that decouples temporally the measurement of muscle activity associated with an LLR–which completes its course within 100 ms of a perturbation–with the measurement of the associated blood-oxygen level dependent (BOLD) signal– which occurs for several seconds after a reflex is elicited (Fig. 1).

**Fig. 1.**
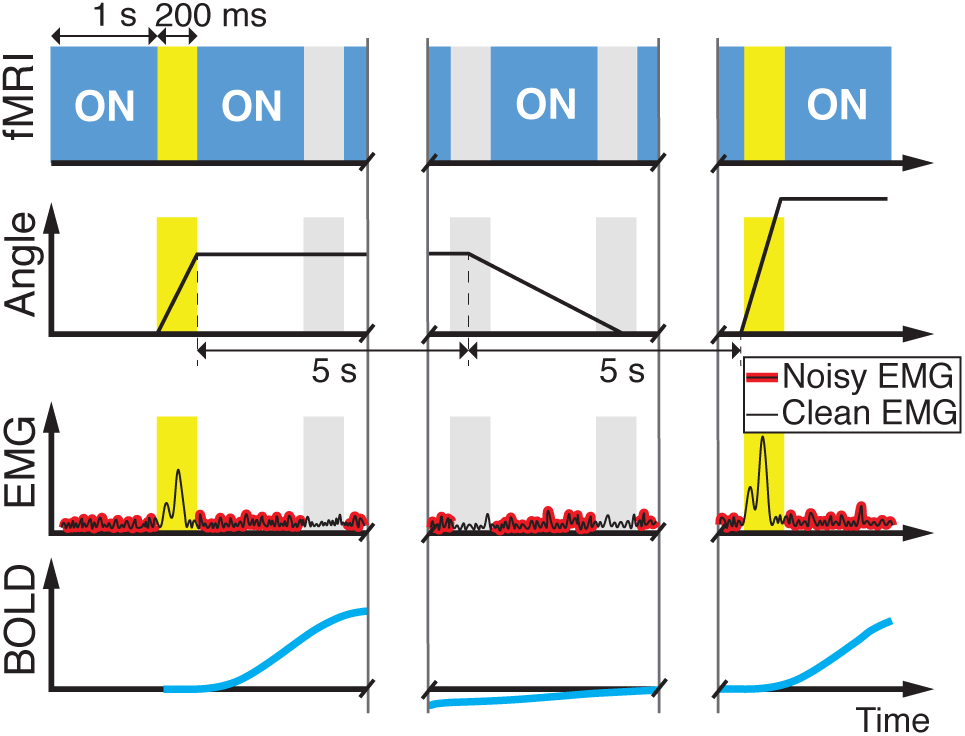
Timing diagram of the MRI sequence (first line) with the 200 ms silent windows between acquisition of different volumes; (second line) commanded robot joint trajectory; (third line) expected EMG signal, which is clean during MRI silent windows; (fourth line) expected BOLD signal associated with the LLR.

While this approach avoids artifacts associated with RF and gradients, preliminary analyses have highlighted the presence of artifacts introduced by the coupling between the static magnetic field and the electrodes movement. As such, we have implemented an optimization-based adaptive filter for the EMG data (see Sec. II-C.2) that determines the set of filtering parameters that minimize the difference between measurements of LLR amplitude taken inside and outside the MRI scanner.

### B. Design of the MR-StretchWrist

Previous studies observed that LLRs can be elicited in the forearm muscles with background wrist torque and perturbation velocities that range from 0 to 0.5 Nm and from 100 to 250 deg/s, respectively. In order to determine the maximum torque characteristic required to apply an effective perturbation, we modeled the wrist joint as a mass-spring system [22], assuming the hand inertia to be *I*_*h*_ = 0.0024 kg m^2^ [22] and considering the stiffness to result from the short-range stiffness behavior of muscle fibers, as modeled in [23]. With these assumptions we determined that to accelerate the hand to 250 deg/s in 50 ms when a background torque of 0.5 Nm is applied, a peak torque *τ*_*R*_ = 3 Nm would be required. To apply such perturbations, we have developed a novel robotic device, the MR-StretchWrist (MR-SW) shown in Fig. 2. The MR-SW is a 1-Degree of Freedom (DOF) robot capable of applying controlled movement to the wrist Flexion/Extension (FE) axis in a range of *θ*_*FE*_ = [−45; 45] deg. It is actuated by a ultrasonic piezoelectric motor (EN60 motor, Shinsei Motor Inc., Japan) which provides 1 Nm peak torque and 900 deg/s peak velocity, previously utilized for several MR-compatible robotic applications [24]–[26]. To fulfill the design specifications, we have employed a capstan transmission with 3:1 gear ratio to transfer motion from the motor to the end effector. The capstan drive consists of two pulleys with different diameters connected together with a smooth cable that, to ensure no-slippage high-friction contact, is wrapped around the pulleys multiple times (Fig. 2). Such transmission is ideal for this application as it is char-acterized by no-backlash, low friction, high-bandwidth properties and it is easily manufacturable using MR-compatible materials. Finally, to measure the wrist joint torque we have instrumented the MR-SW with a six-axis MR-compatible Force/Toque sensor (Mini27Ti, ATI Industrial Automation, Apex, NC).

**Fig. 2.**
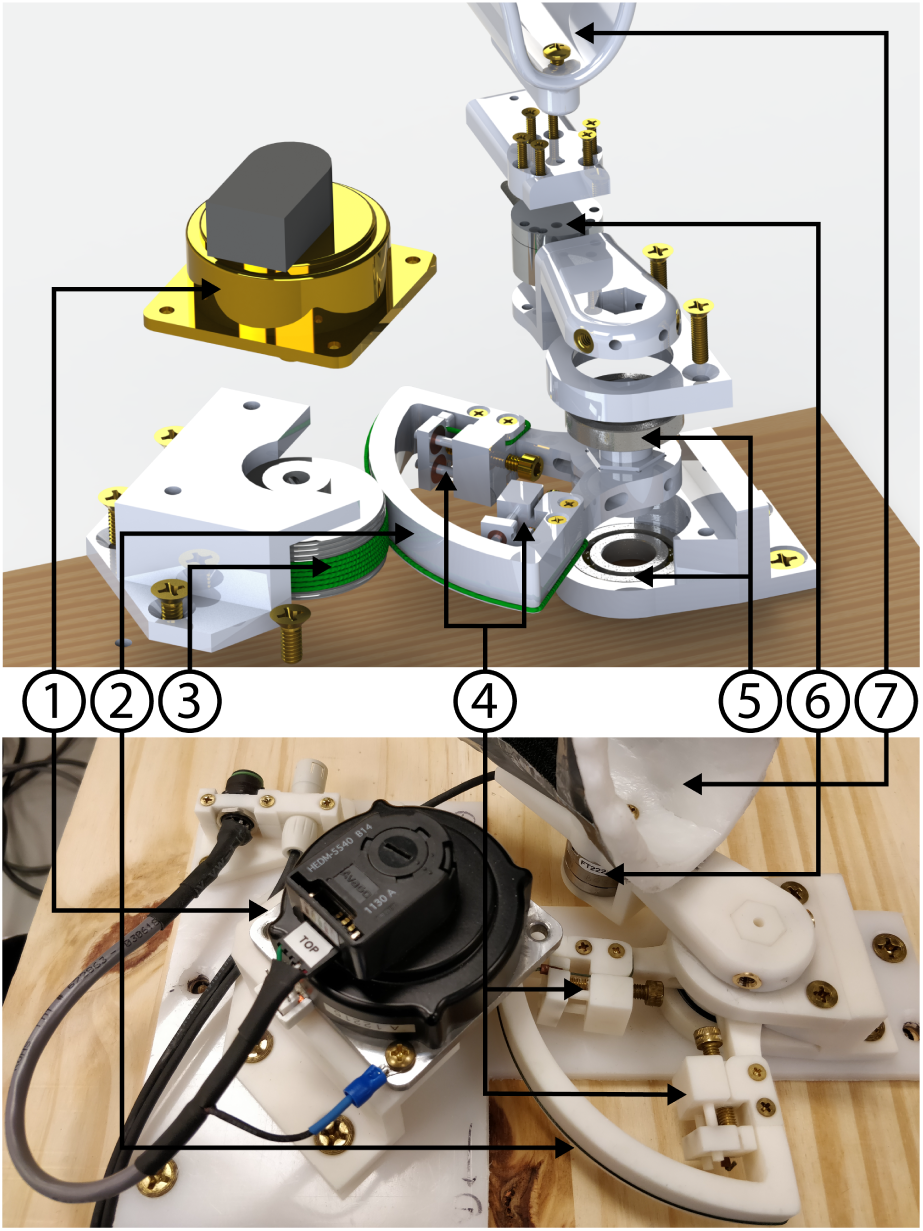
Exploded view (Top) and prototype (Bottom) of the MR-StretchWrist. (1) Ultrasonic motor, (2) Output pulley, (3) Input pulley, (4) Tensioning mechanisms, (5) Structural bearings, (6) Force/Torque sensor, Hand support.

To ensure MR-compatibility of the entire system, all structural components have been manufactured using ABS-based 3D-printed plastic (RS-F2-GPWH-04, Formlabs Inc., MA, USA) and connected using brass screws. For the cable of the capstan transmission we have used a microfiber braided line (SpiderWire Stealth 0.4 mm diameter braided fishing line, 80 lb test). The output shaft is supported by ceramic radial bearings (Boca Bearings, Boynton Beach, FL, USA), while bronze sleeve bearings are included to support the tensioning mechanisms. To reduce the electromagnetic interference introduced in the MRI scanner room by the motor and the motor encoder, a tripolar twisted-pair shielded cable was used for encoder line and shielded cable was used for the motor power line. Both lines are filtered when passing through the scanner patch panel using 5.6 pF and 1.3 pF capacitive filters, respectively.

#### 1) Perturbation protocol

Since the impact dynamics that characterize the end of a perturbation generates undesired oscillations in the EMG recording, each perturbation was kept active for a fixed time of 200 ms, regardless of the value of perturbation velocity. After each perturbation a rest period of 5 s was included before the robot slowly moved the hand back to the neutral position. The return velocity was set to 15 deg/s. A new perturbation was triggered 5 s after the onset of the return movement (Fig. 1). In each of the experimental sessions presented in this paper, we applied perturbations at six different velocities v_des_ =[−200 −150 −100 100 150 200] deg/s when no background torque was applied. Each velocity in v_des_ was repeated 10 times with a pseudo-random order generated for each run. Perturbations have always been applied to the subject’s right hand.

### C. EMG acquisition and processing

Surface EMG was recorded with the BrainVision Recorder software (Brain Products, Munich, Germany) using a 16-channel MR-compatible bipolar amplifier (ExG, Brain Products, Munich, Germany). Given our interest in studying the neural substrates of LLR for both flexors and extensors, the perturbations were applied in both directions, and EMG was recorded from two of the muscles involved in controlling the wrist in flexion and extension: Flexor Carpi Radialis (FCR) and Extensor Carpi Ulnaris (ECU).

#### 1) Data acquisition

After having carefully cleaned the skin with a 70% Isopropyl Alcohol solution, we placed Ag/AgCl bipolar electrodes (multitrode, Brian Products, Munich, Germany) on the muscles belly, parallel to the muscle fibers, with an additional reference electrode that was placed on the lateral epicondyle of the elbow. We then filled the central hole of each electrode with an abrasive Electrolyte-Gel (Abralyt HiCl, Rouge Resolution, Cardiff, UK) to ensure contact between the skin and the electrodes. Contact impedance for each electrode was measured using the BrainVision Recoder software. A cotton swab dipped in abrasive gel was swirled on the skin until the measured contact impedance was lower then 10 kΩ, as described in the product technical specification. In order to minimize movement artifacts, we then carefully taped each electrode and the respective lead on the skin and applied pre-wrap around the entire forearm (Fig. 3).

**Fig. 3.**
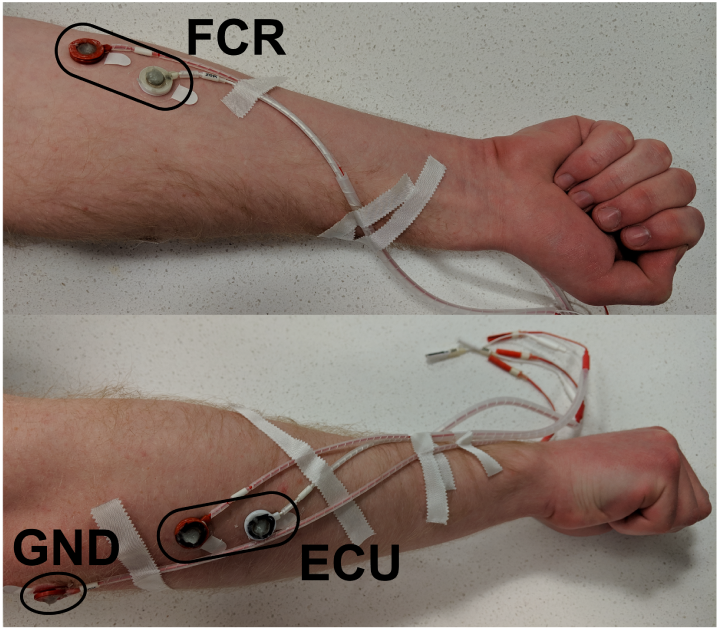
Placement of the electrodes on the forearm

#### 2) Data processing

Pre-processing of the raw EMG data was based on a standard pipeline composed of three steps: band-pass filter to remove motion artifact and high frequency noise (4^th^ order Butterworth filter with cut-off frequencies *f*_*LP*_, and *f*_*HP*_), signal rectification, and low-pass filter to extract the envelope of the EMG signal (4th order Butterworth filter with cut-off frequency *f*_*ENV*_). To extract the magnitude of the long-latency response *H*_*j*_[*i*] that the perturbation *i* elicited on the muscle *j*, we used the *cumsum* method [27], quantifying *H*_*j*_[*i*] as the area underlying the processed EMG signal 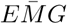 in the temporal window [50, 100] ms after the perturbation onset *t*_0_[*i*]:

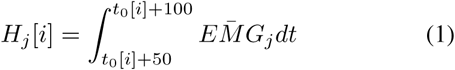

where *t*_0_[*i*] is the time of the *i*-th perturbation onset. Perturbations were classified as reflexive or areflexive by comparison of the metric *H*_*j*_ with a threshold 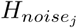 defined as the values that yield 0.05 false positive classification in 50-ms rest periods measured 600 times in the first 30 s of the experimental protocol. To more closely resemble the experimental conditions, the EMG in rest condition was acquired while the MR-SW was applying a position-controlled sinusoidal movement to the wrist joint. The maximum velocity was set to be low enough as to not elicit any reflex (*v*_*max*_ = 20 deg/s). Since the noise characteristic of an EMG recording is intrinsically electrode-specific, we have determined two noise thresholds, one per each EMG channel. For the experimental session performed outside the MRI room (LAB) we set the filtering parameter to be *f*_*LP*_ = 1 kHz, *f*_*HP*_ = 5 Hz, and *f*_*ENV*_ = 100 Hz, while for the MRI experimental condition we determined the filtering parameters **f** *^MRI^* = [*f*_*LP*_, *f*_*HP*_, *f*_*ENV*_] using an optimization-based adaptive filter that seeks to minimize the cost function:

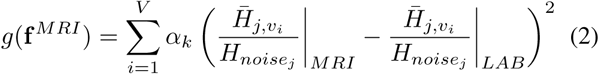

where 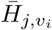 is the average of the metric *H* measured for the muscle *j* when perturbations are applied at the velocity *v*_*i*_; *V* is the number of the different perturbation velocities used in the experimental protocol. *α*_*k*_ is a weight coefficient that accounts for the fact that LLRs are observed when a muscle is both stretched and shorten. We considered *k* = 1 when the perturbation *v*_*i*_ stretches the muscle *j* and *k* = 0 when, instead, perturbation *v*_*i*_ shorten the muscle *j*. Since we expected to have a greater signal-to-noise ratio when measuring LLR of muscles that are being stretched, we set *α*1 = 0.7 and *α*0 = 0.3.

## III Validation Experiments

To validate the different components of the developed methodology, we have performed a set of preliminary experiments by first assessing the MR-compatibility of the MR-SW, and then testing its ability to elicit reflex activity both during MRI scanning and in a laboratory setting. MR-compatibility experiments were conducted at the Center for Biomedical and Brain Imaging c/o University of Delaware using a 64 channel head coil on a Siemens Prisma 3T scanner. Behavioral tests to asses the capability of our methods to elicit and identify a LLR were instead conducted outside the scanner in our lab at University of Delaware on three volunteers.

### A. MR-compatibility

We performed three different experiments to assess the MR-compatibility of the MR-SW. Experiments 1 and 2 were performed as described in [28] to test whether the robotic device: 1. introduces RF noise to the scanner operational frequency band, and 2. increases the temporal signal-to-noise ratio of the MR images acquired using a Multi-Band Accelerated EPI Pulse sequence (CMMR), a standard sequence used for fMRI protocols. Silent acquisition windows were included in the sequence as described in Sec. II-A. Finally, experiment 3 was performed to test whether our experimental protocol introduces signal correlated with the experimental stimulus (i.e. robot motion). This effect would increase the number of false positive voxels identified to correlate with the generation of a LLR, limiting the validity of our results. All test were done using gel phantom (composed of distilled H_2_O doped with 1.25g NiSO_4_x6H_2_O per 1kg H_2_O) that provides temporally and spatially constant signal.

For the methods of experiments 1 and 2 please refer to [28].

For experiment 3 we have used the same acquisition sequence adopted for experiment 2. Seven repeated acquisitions were performed in two different conditions: the first one with the MR-SW outside the scanner room (BL) and the second one with the robot inside the scanner performing a the experimental protocol described in sec. II-B.1 (MVT). Seven additional acquisitions in the BL condition were performed after the removing the robot from the scanner to evaluate the test-retest error of the procedure. To replicate conditions similar to those that characterize our experimental protocol, we positioned the MR-SW and the EMG amplifier at a distance from the scanner’s isocenter that is representative of the one used in nominal operative conditions–i.e. about 1 m away. Moreover, a phantom hand, realized by filling a plastic glove with ager gel, was attached to end effector to resemble the same body mass movement of a human subject experiment.

To quantify the voxel-wise correlation of the image intensity with the three states of the perturbation protocol (REST, PERTURB, RETURN) we ran a first-level analysis using SPM12. A general linear model was constructed using the three states as regressors yielding three F-maps for each of the 21 acquisition. The correlation of the regressor corresponding to the “PERTURB” condition was then quantified as the number of statistically significant voxels in a cuboid ROI placed in the center of the phantom including 50×50×17 = 42500 voxels. Three different uncorrected significance levels were considered *α* = [0.05, 0.01, 0.001] and the number of significant voxels (false positives) was compared between different experimental conditions using two sample t-tests.

### B. Identification of the LLRs

After signing an informed consent, three volunteers have performed a two-session experimental protocol, the first one involving fMRI scanning (MRI) and the second conducted in a lab environment (LAB). During the MRI session subjects were exposed to the protocol described in Sec. II-B.1 while undergoing our modified whole-brain fMRI sequence. The same set of perturbations was then applied in the LAB session that, due to the pseudo-randomic selection of perturbation velocities, was arranged in with different order.

EMG data processing was done as described in the section II-C.2 and the metric *H* was extracted for the FCR and ECU, in both experimental conditions. For each velocity we have then computed the LLR rate, defined as the number of perturbations identified as reflexive over the total number of perturbations, for both LAB and MRI sessions, and calculated the LLR rate error as the difference between the two values, for all muscles, at different velocities.

## IV. Pilot Validation Experiment

One subject was recruited and exposed to an experimental session that featured all the components of the StretchfMRI technique to establish the involvement of RF in the generation of the LLR. The experiment was composed of three sessions, one performed in the lab, necessary to obtain the expected LLR rate required in the adaptive filter, and two performed during fMRI imaging. Imaging parameters included: Multi-Band Accelerated EPI Pulse sequence; 2×2×2 mm^3^ voxel resolution, with 0.3 mm slice spacing, 46 deg degree flip angle, 880×880 px per image, 64×110×110 mm^3^ image volume; TR=1000 ms, and TE=30 ms; pixel band-width=1625 Hz/pixel, receiver gain: high. In each session subjects were exposed to 60 perturbations, pseudo-randomly selected from a pool of six different perturbation velocities ([−200 −150 −100 100 150 200] deg/s) each repeated 10 times, while EMG was recorded from the FCR and ECU muscles.

After the two functional sessions, a high resolution structural scan (magnetization-prepared rapid acquisition with gradient echo (MPRAGE)–1×1×1 mm^3^ resolution, with TR=2300 ms, and TE=3.2 ms; 160 slices with a 256×256 px per image, for a 160×256×256 mm^3^ image volume–was acquired for registration and normalization of the results to a common space. The full experimental protocol took about 85 minutes divided as 20 minutes per each experimental session, 7 minutes for the structural scan and 18 for the robot setup and the electrode placement.

fMRI data was pre-processed in SPM12 using a standard processing pipeline composed of realignment, normalization, and high-pass filtering. We then performed a first-level analysis using the following general linear model:

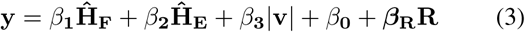

Where 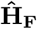 and 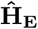 are binary variables that indicate whether a reflex has been triggered during a specific perturbation in the FCR and ECU, respectively. |v| is, instead, a continuous variable that indicate the absolute value of the perturbation velocity. **R** is the regressor of the head movement determined by the data pre-processing. The presence of brain areas whose activity significantly correlates with the generation of a LLR in the FCR and ECU was assessed by analysis of the F-maps corresponding the the coefficients *β*_1_ and *β*_2_, respectively. Significance level was set to p*<* 0.05 uncorrected.

## V. RESULTS

### 1) MR compatibility

Visual inspection of the averaged RF signal intensity spectrum acquired in the MVT condition (Fig. 4 top), showed no peaks of intensity outside the 95% confidence interval, previously determined with 20 repeated measurement in the BL condition, confirming that the MR-SW does not introduce significant RF noise in the scanner.

**Fig. 4.**
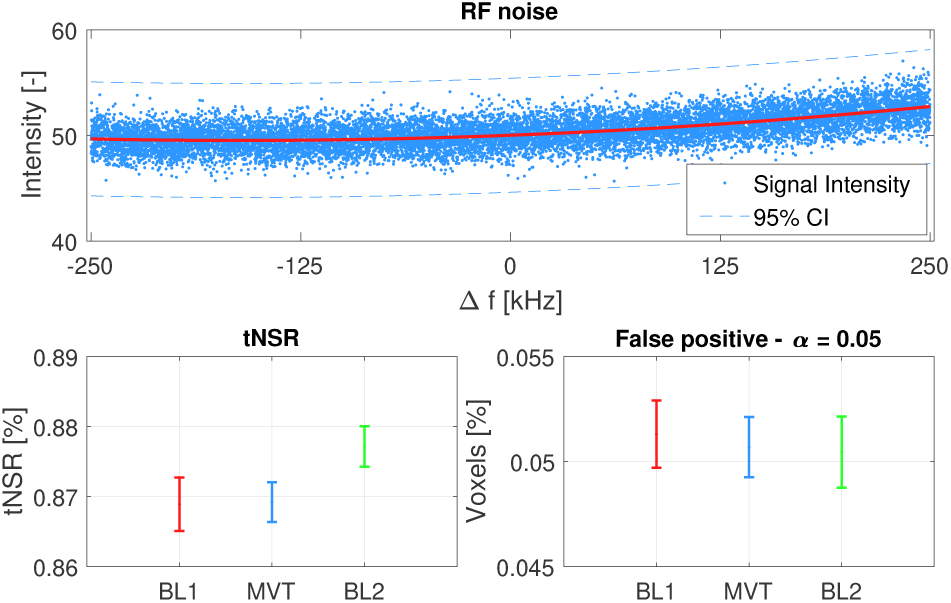
Results of the MR-compatibility experiments. (Top left) 95% confidence intervals of the percent of voxels identified to correlate with the perturbation protocol as a function of the significance level of the first level analysis; (Top right) bootstrapped confidence intervals for mean tNSR for each condition at the p <0.05 level; (Bottom) Frequency spectrum of the RF averaged signal intensity

Distribution of the tNSR values measured in the pre-baseline (BL1), MVT and post-baseline (BL2) conditions are represented in Fig. 4 (bottom left). Pairwise comparisons of the MVT condition with both pre-BL and post-BL conditions fail to reject the null hypothesis that the tNSR measured in the MVT condition is equal to that measured in baseline at a significance level of p *<*0.05, demonstrating that the MR-SW does not significantly alter the tNSR characteristic of the scanner.

Finally, the distributions of the percent number of voxels that significantly correlate (*α* = 0.05) with the application of the velocity controlled perturbations in the three conditions (BL1, MVT, and BL2) are reported in Fig. 4 (bottom right). Two-sample t-test between the MVT and BL conditions fail to reject the null hypothesis that false positive rate is equal in the MVT than in BL condition, for all tested levels of significance. Significance level for the t-test was set to p*<*0.05.

### 2) Identification of the LLRs

Analysis of the EMG signal recorded during LAB experimental sessions shows the capability of our protocol to condition and detect long-latency responses in both FCR and ECU (fig. 5) for all magnitude of applied velocities.

**Fig. 5.**
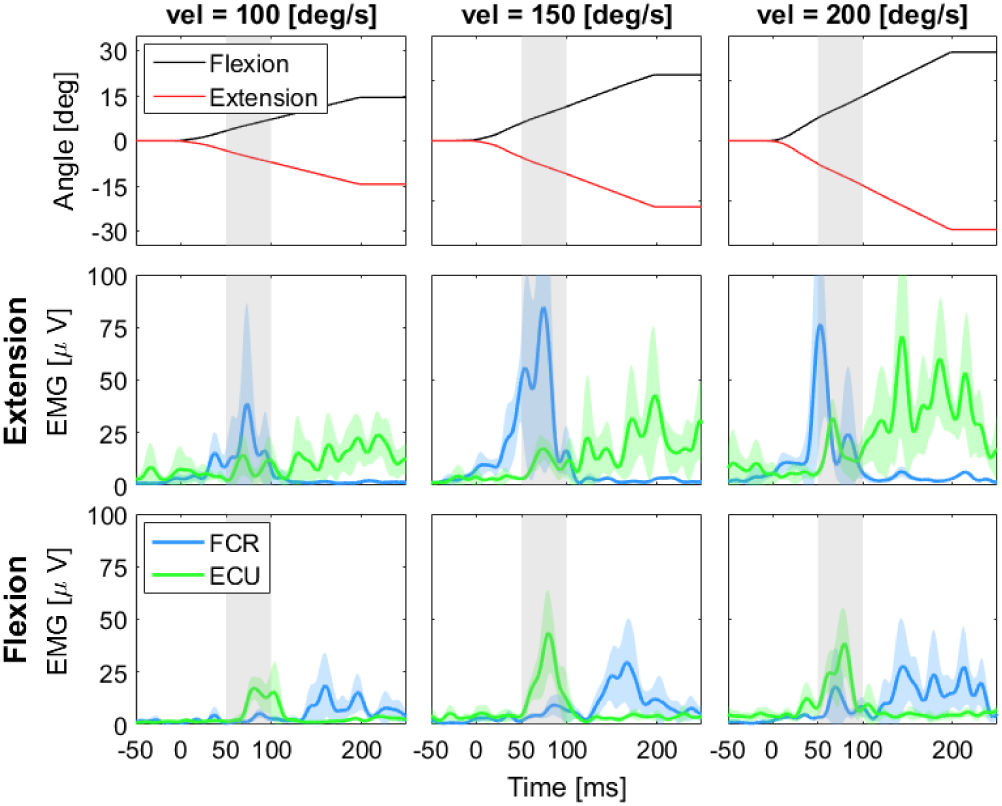
First row: trajectory of the MR-StretchWrist for the different perturbation velocity. Second and third rows: EMG responses measured in a single subject for the three perturbation velocities. In all graph the thick line represents the mean and the shaded area the 95% confidence interval. To facilitate interpretation on the graphs a gray shaded area has been included in each graph in the time interval where a LLR is expected.

**Fig. 6.**
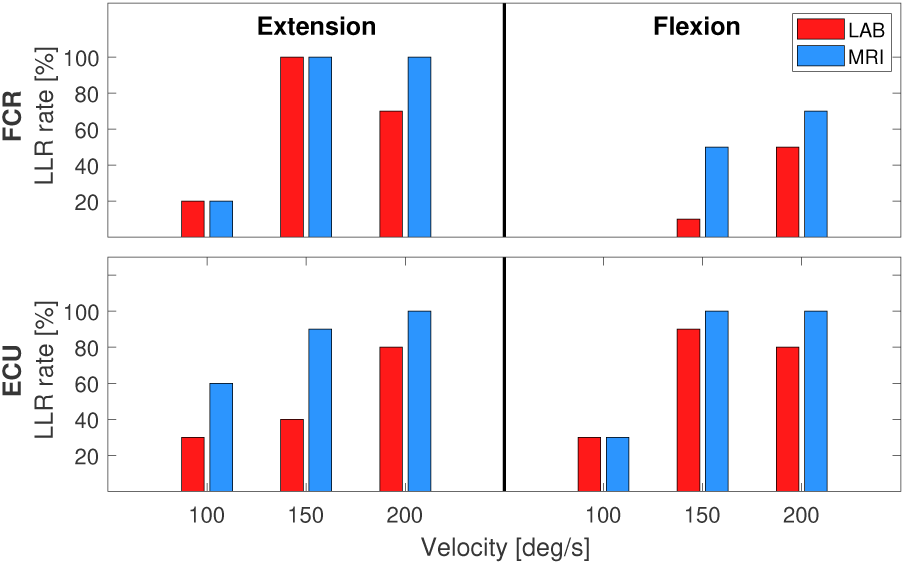
LLR rate measured for the FCR and ECU during sessions done outside the scanner (MRI) and during MRI scanning (MRI)

Finally, comparison between the LLR rate measured in the MRI and LAB conditions shows good agreement for each of the tested values of perturbation velocity (fig.6). The difference between of the LLR rate measured in the MRI and LAB averaged across velocities is 21% in the ECU and 15% in the FCR. Due to the different weighting of the perturbations that stretch and shortens the muscles used in the adaptive filter, for each muscle there is a smaller LLR rate for the direction of perturbation that stretches the muscle compared to the one that shortens it.

#### A. Pilot Experiment

Head movement and rotation were estimated form the imaging parameters and were confirmed to be lower then 1 mm and 1 deg, respectively. Analysis of the activation maps show significant neural activity in the brainstem when LLRs are generated in both FCR and ECU (Fig. 7). The results show a different somatotopical organization for FCR and ECU with the neural activity corresponding to FCR located in the right medullay reticular formation and the activity of the ECU located instead in the left pontine reticular formation.

**Fig. 7.**
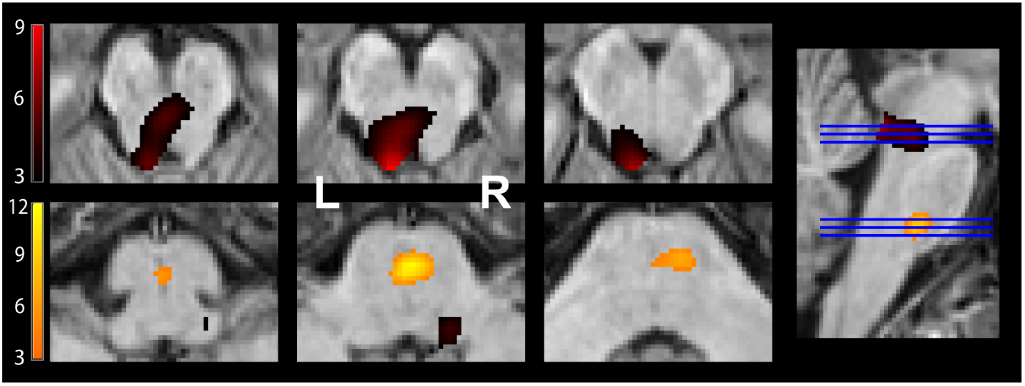
The activation maps that show the neural activity in the brainstem that correlates with the generation of a LLR in the FCR (shown in orange) and ECU (shown in red).

## VI. Discussions and Conclusion

In this work we have presented and validated the StretchfMRI, a novel non-invasive technique that can be used to investigate the neural correlates of the Long-Latency Response. The StretchfMRI technique is composed of three different key elements: an MRI compatible wrist robot (the MR-StretchWrist) capable of safely applying velocity controlled wrist perturbations during fMRI scanning, an MRI sequence that allows for gradient artifact-free recordings of the LLR EMG activity, and an adaptive filtering approach for the EMG data processing that allows for reliable identification of LLRs.

A first set of experiments aimed to test the fMRI compatibility of the experimental protocol, showed that the MR-StretchWrist in operative conditions does not significantly alter the quality of the MR-images, allowing a safe use of the device during fMRI procedures. Analysis of the EMG data recorded in a lab experimental session, where there are no confounders related to the MRI electromagnetic fields, validated our experimental protocol, demonstrated the capability of the MR-SW to elicit LLRs in both flexors and extensors muscles for different levels of applied perturbation velocities. Finally, comparison of the reflex rate measured in lab experiments and during fMRI scanning, that show only an 18% average difference in elicited LLR rate, suggests the capability of our algorithm to reliably identify LLR during fMRI protocol.

A final pilot experiment that integrated all the components of our novel technique showed significant neural activation modulated in the brainstem by the generation of LLRs in both flexors and extensors. The responses of FCR and ECU showed a different somatotopic arrangement as described by the double reciprocal model of the reticulospinal tract, with the neural activity of flexors and extensors located in the medullary and pontine reticular formation, respectively [10].

While a single-subject experiment cannot be used to draw conclusions on the organization of motor function in the reticular formation, our work provided a first preliminary confirmation of the organization of motor responses in the reticular formation previously observed only in animal models. Such encouraging results suggest the StretchfMRI to be a promising tool to investigate the contribution of the RF to motor control and more in general to understand the neural correlates of the processing and generation of long-latency responses.

Our protocol currently used an Echo Planar Imaging (EPI) sequence routinely used for fMRI studies. While the results of our pilot study suggest that our imaging technique has sufficient contrast-to-noise ratio and robustness from motion artifacts to detect meaningful change in deep brainstem nuclei, we are considering to include a high-resolution multishot sequence that has specifically been developed for brainstem imaging expecting a 10-20% increase in contrast-to-noise ratio [29].

## VII. Acknowledgments

We acknowledge support from the University of Delaware Research Foundation grant no. 16A01402, from ACCEL NIGMS IDeA grant no. U54-GM104941, and from startup funds by the University of Delaware.

